# Centrality and the shortest path approach on the human interactome

**DOI:** 10.1101/264069

**Authors:** Natalia Rubanova, Nadya Morozova

## Abstract

A network is one of the most convenient way to represent interactions between biological entities in systems biology. A network of molecular interactions is a graph in which the vertices are biological component and the edges correspond to the interactions between them. Many notions and approaches for network analysis came to systems biology from the theory of graphs – a field of mathematics that study graphs. We focused on the study of the shortest path approach in this work. We investigated whether this approach yields valid molecular paths. To perform this, the shortest paths in the human interactome (derived from HPRD and HIPPIE databases) were found between all relevant combinations of proteins taken from eight well-studied highly conserved signaling pathways from the KEGG database (NF-kappa B, MAPK, Jak-STAT, mTOR, ErbB, Wnt, TGF-beta and the signaling part of the apoptotic process). Canonical paths were systematically compared with the shortest counterparts and centrality of vertices and paths in subnetworks induced by the shortest paths were analyzed. We found that the sets of the shortest paths contain the canonical counterparts only for very short canonical paths (length 2-3 interactions). We also found that high centrality vertices tend to belong to canonical counterparts, to less extent this can also be said about high centrality paths.

## Introduction

Modern molecular biology accumulated vast amount of knowledge about interactions between molecular entities within the cell. This knowledge is stored in databases of molecular interactions such as HPRD (Peri et al. 2003), Hippie (Schaefer et al. 2012), bioGrid (Stark 2006), STRING (Jensen et al. 2009). Networks are the basic tool to visualize and to analyze these interactions (Kitano 2002; Barabási & Oltvai 2004). A network of molecular interactions is a direct or an indirect graph where vertices represent biological entities (e.g. proteins, genes, small molecules, metabolites, …) and edges represent interactions between them such as physical interactions, gene regulation, chemical reaction.

Numerous concepts for network analysis came to systems biology from the graph theory – a field of mathematics that studies graphs. These are the concepts such as degree of a vertex (Pavlopoulos et al. 2011) – which is the number of edges of a vertex; degree distribution – probability distribution of vertex degree over the whole network (Pavlopoulos et al. 2011); betweenness, closeness, eigenvector, degree centrality – different measures of importance of a vertex in a network (Pavlopoulos et al. 2011); clustering coefficient – the measurement that shows the tendency of a graph to be divided into clusters (Pavlopoulos et al. 2011); the shortest path between two vertices – such a path that the sum of weights or the length is minimum possible in the network (Pavlopoulos et al. 2011).

The applications of these concepts greatly vary. Vertex degree and centrality is used to distinguish possible hubs – vertices that might be important for biological system (Milioli et al. 2017; Dong et al. 2016). Clustering coefficient – to analyze network topology (Hao et al. 2012). The shortest path approach is used to construct networks for a set of genes (Yuan et al. 2017), to prioritize vertices (Zhang et al. 2012), to predict functional components and molecular pathways (Nakamura et al.2012; Bromberg et al. 2009), to perform network modularization (Cabusora et al. 2005) and to predict protein function (Sharan et al. 2007).

In this work, we studied whether the shortest path approach when used alone on human interactome can yield valid molecular paths and whether high centrality vertices and paths in the subnetworks induced by the shortest paths belong to canonical counterparts. The reasoning behind using centrality is the following. Centrality measures were first introduced for social network analysis to identify ‘important’ persons in a given social network. Several types of the centrality measure were developed depending on what the ‘important’ could mean (betweenness, closeness, eigenvector, degree centrality). Recent studies also show that high centrality vertices can be related to network stability. Our goal was to check whether ‘important’ vertices from the network point of view would be from the canonical pathways.

To answer the abovementioned questions, we compared linear paths (we name them canonical paths) taken between all relevant pairs of source/target point on 8 canonical pathways from the KEGG database with the shortest path counterparts and checked whether high centrality vertices and paths in the subnetworks induced by the shortest paths belong to canonical paths. The following pathways were used: NF-kappa B, MAPK, Jak-STAT, mTOR, ErbB, Wnt, TGF-beta signaling pathways and the signaling part of the apoptotic process. These are well-studied highly conserved signaling pathways that regulate a wide range of cellular processes such as proliferation, differentiation, inflammation, apoptosis, angiogenesis, adhesion and migration. Importantly, the signal transduction mechanism of the pathways works via protein-protein interactions.

The pathways were dissected into linear paths and all relevant combinations between source and target points were taken for the analysis. This resulted in 91 paths of length 2 interactions, 74 paths of length 3 interactions, 51 pathways of length 4 interactions and 29 pathways of length 5 interactions. The shortest paths between source and target points of the paths were taken in four networks created using protein-protein interaction from HPRD database, HIPPIE database with only high confidence interactions, HIPPIE database with high and medium confidence interactions, HIPPIE with high, medium and low confidence interactions. The shortest paths as well as high centrality vertices and paths in subnetworks induced by the shortest paths were compared with canonical counterparts. We found that one of the shortest paths match the canonical counterpart for 56-77% of the canonical paths of length 2 interactions. One of the shortest paths match the canonical counterpart only for 15-30% of the canonical paths of length 3 interactions. None of the shortest paths match the canonical counterpart for canonical paths of length 4 and 5 interactions. However, we found that the vertices with high centrality scores belong to canonical pathways for *45-75%* of the paths depending on the length and the database. The paths with high centrality score belong to canonical pathways for *17-54%* of the paths also depending on the length and the database.

## Materials and Methods

### Databases

The following databases were used to create protein-protein networks: Human Protein Reactions Database (HPRD) v9.1 and Human Integrated Protein-Protein Interaction rEference (HIPPIE) v2.0. Both databases contain experimentally validated protein-protein interactions for human cells. The confidence score is assigned for each interaction in HIPPIE database. The confidence score is calculated as a weighted sum of the number of studies in which an interaction was detected, the number and quality of experimental techniques used to measure an interaction and the number of non-human organisms in which an interaction was reproduced (Anon n.d.). We used predefined confidence levels by HIPPIE team to create networks with only High Confidence interactions – HIPPIE HC (confidence level 0.73 – third quartile of the HIPPIE score) and with High and Medium Confidence interactions – HIPPIE HC+MC (confidence level 0.63 – second quartile of the score distribution).

### Pathways

The parts of NF-kappa B, MAPK, Jak-STAT, mTOR, ErbB, Wnt, TGF-beta signaling pathways, the signaling part of the apoptotic process consisting only of protein-protein interactions were taken from KEGG database(Ogata et al. 1999) for the analysis. 55 simple linear protein-protein paths of length from 2 to 5 interactions were found for 8 pathways (Supplementary Table 1). All possible combinations of source and target points preserving the direction of signal transduction were found from these 55 paths that gave 238 paths of the length from 2 to 5 interactions: 91 paths of length 2 interactions, 74 paths of length 3 interactions, 51 pathways of length 4 interactions and 29 pathways of length 5 interactions (Supplementary Table 2).

**Table 1.**
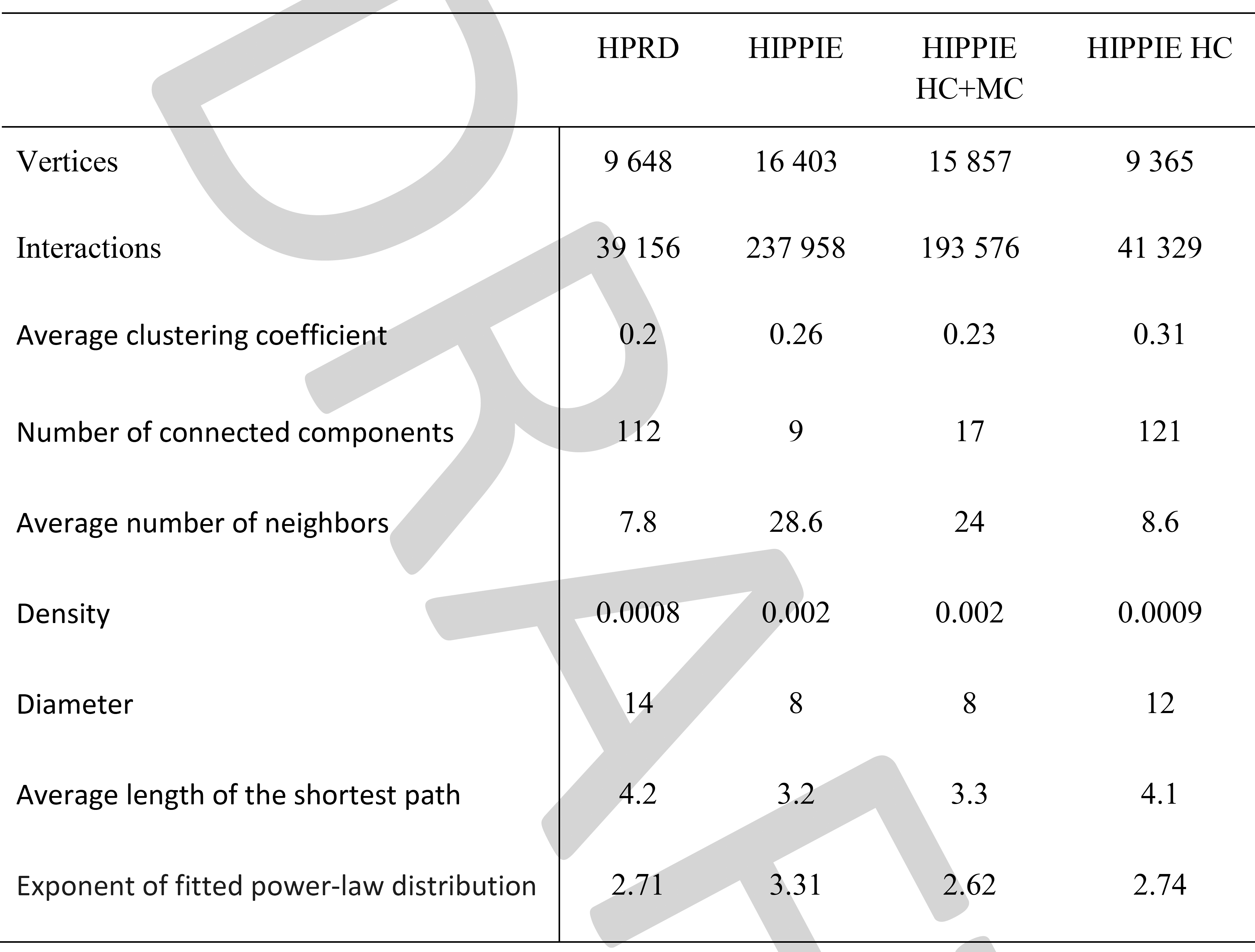
Basic topological properties of the networks created using HPRD and HIPPIE database.

### The shortest paths

The shortest paths between each pair of source and target points in each 4 protein-protein networks were found using the breadth-first search algorithm. In such cases when a canonical path starts and/or ends from a protein complex each subunit of the complex were taken as a source/target points. This procedure results in a set of the shortest paths for every pair of source/target points. Every path in the set was compared with the canonical counterpart.

### Construction of subnetworks of the shortest paths and centrality scores

Networks for each pair of source/target points were created from the set of the shortest paths without duplicated edges. Centrality scores were calculated for each vertex and path except the source/target points as number of the shortest paths that pass through the vertex or the path. Vertices with centrality scores higher than the upper quartile for a network and paths with centrality scores higher than average for a network were compared with canonical counterparts.

## Results

Table 1 gives information about basic topological properties for the created networks. The exponent of the fitted power-law distribution in the degree distribution was calculated with powerlaw (Alstott et al. 2014) Python package. It can be seen, that the topological properties of the networks created from HPRD and HIPPIE HC (only high confidence interactions) databases drastically differs from the topological properties of the networks created from HIPPIE and HIPPIE HC+MC (only high and medium confidence interactions) databases.

We compared the lengths of the shortest paths and the lengths of the canonical counterparts. We found that the average length of the shortest paths is less by at least 1 interaction for the canonical paths of length 4 and 5 interactions (Figure 1) and is still less than the average length of the shortest path in the networks.

**Figure 1.**
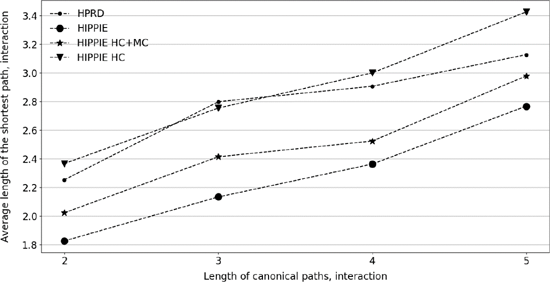
Average length of the shortest paths.

Next, we checked whether one of the paths in the set of the shortest paths will match the canonical counterpart. We found that the set of the shortest paths contains the canonical counterpart only for 56-77% of the canonical paths of length 2 interactions, for 15-30% of the canonical paths of length 3 interactions and for 0% of the canonical paths of length 4 and 5 interactions (Figure 2).

**Figure 2.**
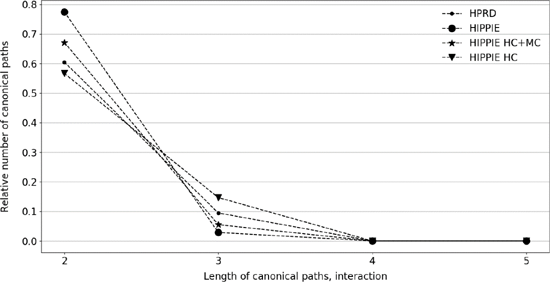
Relative number of canonical paths for which the sets of the shortest paths contains the canonical counterpart.

We also checked whether vertices and paths with high centrality scores belong to the canonical counterpart. We found that the vertices with centrality scores higher than the upper quartile belong to the canonical counterpart for 46-77% of the canonical paths depending on database (Figure 3a). The paths with centrality more than average belong to the canonical counterpart only for 16-54% paths (Figure 3b)

**Figure 3.**
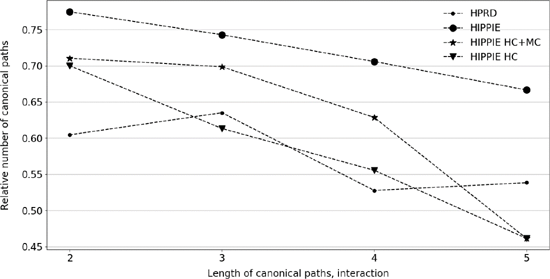
Relative number of canonical paths for which high centrality vertices in subnetworks induced by the shortest paths belong to the canonical counterpart

**Figure 4.**
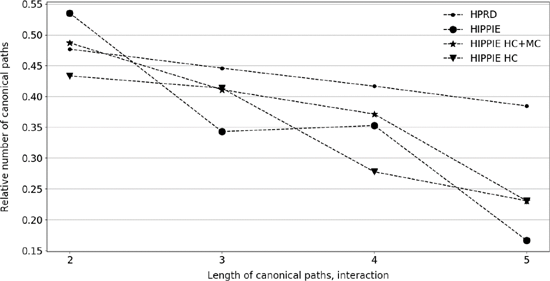
Relative number of the canonical paths for which high centrality paths in subnetworks induced by the shortest paths belong to the canonical counterpart.

## Discussion and Conclusion

The shortest path approach is widely used approach to find connections between vertices in systems biology. We studied in this paper whether such an approach could yield valid paths.

We performed an experiment by comparing the shortest paths between all possible source and target points on eight canonical pathways with the canonical counterparts. We found that the shortest paths between these points built in the human interactome constructed from HPRD or HIPPIE databases usually do not match the canonical counterpart. We also examined whether the vertices and the paths with high centrality score in the network created using the shortest path between these points belong to the canonical counterpart. We found that the vertices with centrality score higher than the upper quartile belong to the canonical counterpart for up to 77% of the canonical paths and the paths with centrality more than average belong to the canonical counterpart for up to 54% of the canonical paths.

It is interesting that the network constructed from the whole HIPPIE database shows completely opposite performance for vertices and paths with high centrality score. A probable explanation for this is that many valid protein-protein interactions are assigned low confidence score, but on the other hand low confidence interactions that include false positives interactions could introduce shortcuts into the shortest paths.

We conclude that the shortest path between two vertices in the human interactome usually does not match the valid biological path. However, studying additional topological features like centrality of a vertex or of a path in the network constructed from the shortest paths might help at least partly reconstruct the valid biological path.

## Supplementary Table 1. List of paths

**NFKB:**

TNFRSF11A-TRAF6, TRAF2-MAP3K14-CHUK-NFKB2, RELB;

TNFRSF11A-TRAF6, TRAF2-TAB1, TAB2, TAB3-IKBKG, CHUK, IKBKB-NFKBIA;

LTBR-TRAF2, TRAF5-MAP3K14-TAB1, TAB2, TAB3, MAP3K14-IKBKG, CHUK, IKBKB-NFKBIA; LTBR-TRAF2, TRAF3-MAP3K14-CHUK-NFKB2, RELB;

TNFRSF13C-TRAF2, TRAF3-MAP3K14-CHUK-NFKB2, RELB;

CD40-TRAF2, TRAF3-MAP3K14-CHUK-NFKB2, RELB;

CD40-TRAF6-TAB1, TAB2, TAB3-IKBKG, CHUK, IKBKB-NFKBIA; TLR4-TICAM2, TICAM1-RID1, TRAF6-TAB1, TAB2, TAB3, MAP3K14-IKBKG, CHUK, IKBKB-NFKBIA;

TLR4-TIRAP, MYD88-IRAK1, IRAK4, TRAF6-TAB1, TAB2, TAB3-IKBKG, CHUK, IKBKB-NFKBIA; DDX58-TRAF2, TRAF6-TAB1, TAB2, TAB3, MAP3K14-IKBKG, CHUK, IKBKB-NFKBIA; TNFRSF1A-RIPK1, TRADD, TRAF2, TRAF5-TAB1, TAB2, TAB3, MAP3K7-IKBKG, CHUK, IKBKB-NFKBIA;

IL1R1-MYD88, IRAK1, IRAK4, TRAF6-TAB1, TAB2, TAB3, MAP3K7-IKBKG, CHUK, IKBKB-NFKBIA;

**MAPK:**

NTRK1, NTRK2-GRB2-SOS1, SOS2, RRAS2, MRAS, HRAS, KRAS, NRAS, RRAS-BRAF, RAF1, MOS-MAP2K1, MAP2K2;

EGFR-GRB2-SOS1, SOS2, RRAS2, MRAS, HRAS, KRAS, NRAS, RRAS-BRAF, RAF1, MOS-MAP2K1, MAP2K2;

FGFR1, FGFR2, FGFR3, FGFR4-GRB2-SOS1, SOS2, RRAS2, MRAS, HRAS, KRAS, NRAS, RRAS-BRAF, RAF1, MOS-MAP2K1, MAP2K2;

PDGFRA, PDGFRB-GRB2-SOS1, SOS2, RRAS2, MRAS, HRAS, KRAS, NRAS, RRAS-BRAF, RAF1, MOS-MAP2K1, MAP2K2;

TNFRSF1A, ILR1, ILR2-CASP3-PAK1, PAK2=MAP3K1-MAP2K4-MAPK8, MAPK9, MAPK10; TNFRSF1A, ILR1, ILR2-TRAF2-MAP3K5-MAP2K3, MAP2K6-MAPK14, MAPK11, MAPK13, MAPK12; TNFRSF1A, ILR1, ILR2-TRAF6-MAP3K7-NLK;

TNFRSF1A, ILR1, ILR2-TAB1-MAP3K7-NLK;

**JAKSTAT:**

IL22RA2, CNTFR, CSF2RA, CSF2RB, CSF3R, IL23R, IFNLR1, EPOR, GHR, IFNAR2, IFNGR1, IFNGR2, IFNGR2, IL2RA, IL2RB, IL2RG, IL3RA, IL4R, IL5RA, IL6R, IL6ST, IL7R, IL9R, IL10RA, IL10RB, IL11RA, IL12RB1, IL2RB2, IL13RA1, IL13RA2, IL15RA, LEPR, LIFR, MPL, IL21R, IL20RA, IL20RB, PRLR, IL22RA1, CRLF2, OSMR, IL27RA-PTPN11, GRB2-SOS1, SOS2-HRAS-RAF1;

**WNT:**

WNT16, WNT4, WNT1, WNT2, WNT3, WNT5A, WNT6, WNT7A, WNT7B, WNT8A, WNT8B, WNT10B, WNT11, WNT2B, WNT9A, WNT9B, WNT10A, WNT5B, WNT3A-FZD10, FRZD2, FZD5, FZD3, FZD4, FZD6, FZD7, FZD8, FZD9, LRP6, LRP5-DVL1, DVL2, DVL3-GSK3B-CTNNB1-LEF1, TCF7, TCF7L2, TCF7L1;

**ERB:**

EGFR-SRC-PTK2;

EGFR-CRK, CRKL-ABL1, ABL2;

EGFR-NCK1, NCK2-PAK4, PAK1, PAK2, PAK3, PAK6, PAK5-MAP2K7, MAP2K4-MAPK8, MAPK9, MAPK10-JUN, ELK1;

ERBB2-SRC-PTK2;

ERBB2-CRK, CRKL-ABL1, ABL2;

ERBB2-NCK1, NCK2-PAK4, PAK1, PAK2, PAK3, PAK6, PAK5-MAP2K7, MAP2K4-MAPK8, MAPK9, MAPK10-JUN, ELK1;

**MTOR:**

WNT16, WNT4, WNT1, WNT2, WNT3, WNT5A, WNT6, WNT7A, WNT7B, WNT8A, WNT8B, WNT10B, WNT11, WNT2B, WNT9A, WNT9B, WNT10A, WNT5B, WNT3A-DVL1, DVL2, DVL3 -GSK3B-TSC1, TSC2, TBC1D7-RHEB;

TNF-TNFRSF1A-IKBKB-TSC1, TSC2, TBC1D7-RHEB;

IGF1, INS-IGF1R, INSR-IRS1-PIK3CA, PIK3CB, PIK3D, PIK3R1, PIK3R2, PIK3R3;

**APOPTOSIS:**

TNFRSF10D, TNFRSF10C, TNFRSF10B, TNFRSF10A, FAS-FADD-CASP8, CASP10-CASP3, CASP7-TUBA1B, TUBA3E, TUBA3D, TUBA4A, TUBA8, TUBA3C, TUBA1A, TUBAL3, TUBA1C, MCL1, ACTB, ACTG1, SPTA1, SPTAN1, LMNA, LMNB1, LMNB2, PARP2, PARP3, PARP1, PARP4, DFFA, DFFB;

TNFRSF1A-FADD, TRADD-CASP8-CASP3-TUBA1B, TUBA3E, TUBA3D, TUBA4A, TUBA8,

TUBA3C, TUBA1A, TUBAL3, TUBA1C, MCL1, ACTB, ACTG1, SPTA1, SPTAN1, LMNA, LMNB1, LMNB2, PARP2, PARP3, PARP1, PARP4, DFFA, DFFB;

TNFRSF1A-FADD, TRADD-CASP8-CASP7-TUBA1B, TUBA3E, TUBA3D, TUBA4A, TUBA8,

TUBA3C, TUBA1A, TUBAL3, TUBA1C, MCL1, ACTB, ACTG1, SPTA1, SPTAN1, LMNA, LMNB1, LMNB2, PARP2, PARP3, PARP1, PARP4, DFFA, DFFB;

TNFRSF1A-FADD, TRADD-CASP10-CASP3-TUBA1B, TUBA3E, TUBA3D, TUBA4A, TUBA8, TUBA3C, TUBA1A, TUBAL3, TUBA1C, MCL1, ACTB, ACTG1, SPTA1, SPTAN1, LMNA, LMNB1, LMNB2, PARP2, PARP3, PARP1, PARP4, DFFA, DFFB;

TNFRSF1A-FADD, TRADD-CASP10-CASP7-TUBA1B, TUBA3E, TUBA3D, TUBA4A, TUBA8, TUBA3C, TUBA1A, TUBAL3, TUBA1C, MCL1, ACTB, ACTG1, SPTA1, SPTAN1, LMNA, LMNB1, LMNB2, PARP2, PARP3, PARP1, PARP4, DFFA, DFFB;

TNFRSF1A-TRADD, RIPK1, TRAF-DAB2IP-MAP3K5-MAPK8, MAPK9, MAPK10-JUN, FOS;

FAS-DAXX-MAP3K5-MAPK8, MAPK9, MAPK10-JUN, FOS;

ERN1, TRAF2-MAP3K5-MAPK8, MAPK9, MAPK10-JUN, FOS;

CTSC, CTSB, CTSD, CTSH, CTSK, CTSL, CTSV, CTSO, CTSS, CTSW, CTSZ, CTSF-BIRC2, BIRC3, BIRC5, XIAP-CASP9-CASP7, CASP6-TUBA1B, TUBA3E, TUBA3D, TUBA4A, TUBA8, TUBA3C, TUBA1A, TUBAL3, TUBA1C, MCL1, ACTB, ACTG1, SPTA1, SPTAN1, LMNA, LMNB1, LMNB2, PARP2, PARP3, PARP1, PARP4, DFFA, DFFB;

